# Deciphering the role of intrinsically disordered regions of FUS in recognizing U1 snRNA

**DOI:** 10.64898/2026.06.03.729848

**Authors:** Swarnadip Mitra, Farindra Kumar Mahto, Atanu Maity, Ranjit Prasad Bahadur

## Abstract

Fused in Sarcoma (FUS) is an RNA-binding protein associated with neurodegenerative disorders such as amyotrophic lateral sclerosis and frontotemporal dementia. Along with its structured RNA recognition motif, FUS contains two intrinsically disordered regions (IDRs) that play important roles in RNA recognition. However, structural mechanism and dynamics of these IDRs in recognizing RNA remain elusive. We have used molecular dynamics simulations to investigate the structure and dynamics of the two IDRs, flanking the RRM domain of human FUS, in both Apo and U1 snRNA-bound states. Comparison of structural parameters and molecular interactions reveals that RNA binding stabilizes the IDRs and reduces their conformational flexibility through reorganization of intra-protein contacts and correlated movements. RNA binding also causes rearrangement of backbone dihedral angles of the IDRs and limits the formation of secondary structures such as α-helices and 3_10_ helices. Interestingly, the two IDRs exhibit distinct modes of RNA recognition. The N-terminal IDR interacts mainly with the nucleotides located near the terminal regions of the stem-loop snRNA. On the other hand, the C-terminal IDR preferentially associates with the central double-stranded region through extensive interactions with the minor groove of the snRNA. Even within IDRs, we identify specific residues that associate with the snRNA more frequently than others by forming persistent hydrogen bonds. Overall, our findings suggest that the flanking disordered regions of FUS are not merely flexible linkers, but actively participate in RNA recognition through distinct interaction mechanisms. These results provide molecular-level insights into the functional roles of IDRs in recognizing stem-loop RNA.

## 1. Introduction

Intrinsically disordered proteins (IDPs) contain unfolded flexible regions, defined as intrinsically disordered regions (IDRs), which are stable under physiological conditions in cell [1]. IDRs provide the necessary flexibility to acquire a wide range of conformational polymorphism [2-4], which enables IDPs to bind various partner molecules and participate in diverse molecular functions [5, 6]. IDPs exist in an ensemble of conformations ranging from fully ordered to fully disordered with intermediate mixed conformations [7]. IDRs frequently contain transitional elements like Molecular Recognition Features (MoRFs), Short Linear Motifs (SLiMs) and Low Complexity Regions (LCRs), which undergo disorder-to-order transition upon binding their targets [5,7]. Besides binding to other proteins, IDPs frequently bind DNA and RNA [8]. RNA binding proteins (RBPs) are enriched with IDRs and are crucial biomolecules as they regulate key processes of central dogma including transcription, mRNA splicing and editing, translation and post translational modifications (PTMs) [4,9,10]. Several RBPs including transcription factors (TFs) [11], phosphatase [12], phosphorylase [13], nuclease [14] and RNA polymerase [15] contain disordered regions. Several IDPs in humans are also involved in various diseases [16,17], including cystic fibrosis [18], diabetes [19] and various cancer pathways [16,20]. Several RNA binding IDPs play key roles in neurodegenerative disorders [21] including Alzheimer’s Disease (AD) [22,23], Parkinson’s Disease (PD) [24,25], Frontotemporal Dementia (FTD) [26] and Amyotrophic Lateral Sclerosis (ALS) [27,28].

Fused in Sarcoma protein (FUS) is an RNA-binding protein containing both structured domains and intrinsically disordered regions, and is implicated in human neurodegenerative diseases such as amyotrophic lateral sclerosis (ALS) and frontotemporal dementia (FTD) [28]. Human FUS is a 526-residue protein involved in diverse cellular processes, including RNA splicing, RNA transport and DNA repair [29]. Mutations in FUS promote aberrant cytoplasmic localization and aggregation of proteins, which are associated with ALS and FTD [30-32]. FUS contains an RNA recognition motif (RRM) spanning residues from 285 to 371 as annotated by UniProt [33]. FUS interacts with snRNA through multiple regions, including Arginine-Glycin-Glycin (RGG) motif, zinc finger (ZnF) domain and low complexity domain (LCD) [34]. The RRM domain of FUS binds with the stem-loop RNA structure. FUS has two IDRs, which flank the RRM domain. The N-terminal IDR (nIDR) spans from residues 260 to 287, while the C-terminal IDR (cIDR) spans from residues 375 to 390. The nIDR overlaps with the RGG1 motif, and the cIDR overlaps with the RGG2 motif. Structural studies of the FUS-RRM bound to stem-loop RNA revealed additional interactions involving residues flanking the RRM domain [35]. These residues belong to RGG1 and RGG2 regions, which exhibit substantial intrinsic disorder. Conformational heterogeneity and the dynamic nature of these disordered regions contribute to transient and multivalent interactions with RNA, making their structural characterization challenging. This heterogeneity is partly reflected in the ensemble of NMR structures. However, the NMR ensemble is still limited compared to the extensive conformational landscape accessible to IDRs. Computational modelling and molecular simulation can therefore complement experimental approaches by exploring the conformational dynamics of these IDRs in greater detail.

In this study, we have carried out molecular dynamics (MD) simulations of FUS RRM bound to U1 snRNA stem-loop to understand the role of its flanking disordered regions in RNA binding. A comparison between the Apo and the RNA-bound states shows that RNA binding stabilizes both the nIDR and the cIDR, and reduces the overall conformational flexibility of the protein. Interestingly, the nIDR and the cIDR exhibit distinct conformational dynamics, particularly in terms of structural fluctuations and transient secondary structure formation. Analyses of hydrogen bonds (H-bond) and solvent accessibility further reveal highly dynamic interactions between the snRNA and the IDRs. Overall, our results suggest that these RNA-binding IDRs in FUS are not merely flexible linkers, but also contribute differently to recognize snRNA.

## 2. Materials and Methods

### 2.1. Selection of disordered regions and identification of interacting regions

Three-dimensional (3D) NMR structure of the RRM domain of FUS bound to U1 snRNA (PDB id: 6SNJ) [36] was retrieved from the Protein Data Bank (PDB) [37]. Annotation of secondary structures of the FUS RRM was taken from the PDB. Disorder probability score for each residue was obtained from the PDB, which uses IUPred2-short [38] disorder predictor to calculate disorder probability score. The stretch of residues from 260 to 287 at the N-terminal of FUS was defined as nIDR, whereas the stretch of residues from 375 to 390 at the C-terminal was defined as cIDR. We identified RRM residues of FUS interacting with the nucleotides of U1 snRNA, if any atom pair between them are within a distance cutoff of 4.5 Å. Solvent accessible surface area (SASA) of the protein-RNA complex, as well as that of the unbound protein and the unbound RNA were calculated using the program NACCESS [39]. Buried surface area (BSA) contributed by FUS at the protein-RNA interface is calculated by subtracting the SASA of the bound FUS from that of the unbound FUS. Similarly, we also calculated BSA contributed by the snRNA. We also estimated BSA contributed by the nIDR and the cIDR to the overall complex interface.

### 2.2. System preparation for the MD simulation

In this study, we have used the NMR structure of FUS bound to U1 snRNA (PDB ID: 6SNJ). The unbound FUS is prepared by extracting the coordinates of FUS from the complex structure and is referred to as Apo. Each system was solvated in a transferable intermolecular potential with TIP3P water box with a minimum thickness of 1.2 nm from the protein surface to the edge of the box. The final box dimensions were 8.26 ×8.26 ×5.84 nm^3^. The systems were neutralized by adding *Na*^+^and *Cl*^−^ ions at random positions to achieve a physiological salt concentration of 0.15 M.

### 2.3. Molecular dynamics simulation of bound and unbound FUS

All-atom MD simulations were performed using GROMACS 2025.3 [40,41] with the AMBER14 [42,43] force field parameters. Periodic boundary conditions were applied in all three dimensions to eliminate edge effects. A cut-off distance of 1.2 nm was used for short-range van der Waals interactions, while long-range electrostatic interactions were treated using the Particle Mesh Ewald (PME) [44] method. The systems were solvated with pre-equilibrated TIP3P water molecules consistent with the selected force field. Bond lengths involving hydrogen atoms in the protein were constrained using the LINCS [45] algorithm, and water geometry was maintained using the SETTLE [46] algorithm with an integration time step of 2 fs. Each system was energy minimized using steepest descent algorithm to remove steric clashes. This was followed by equilibration in the NVT ensemble at 300 K for at least 5 ns with temperature controlled using the modified Berendsen thermostat (V-rescale) [47]. Post-equilibration, production simulations were carried out in the NPT ensemble at 300 K and 1.0 bar pressure. Pressure was regulated using the Parrinello–Rahman barostat [48], ensuring stable density and box dimensions. Trajectory coordinates were saved every 10 ps for further analysis. A single 1.8 μs trajectory was generated for the Apo system. For the RNA-bound complex, two independent trajectories of equal length of 1.8 μs each were generated and termed as Traj1 and Traj2. Along with the Apo system, this resulted in a cumulative simulation time of 5.4 μs.

### 2.4. Principal Component Analysis (PCA)

MD simulations generate large datasets, making analysis challenging when considering the entire system. PCA is a powerful technique used for dimensionality reduction without losing significant information [49,50]. It helps to identify the dominant motions within the system. For protein, PCA captures the major collective motions that contribute most to the overall dynamics [51]. First, overall rotational and translational motions were removed from the trajectories. Then, the covariance matrix of atomic fluctuations is constructed and diagonalized to obtain orthogonal eigenvectors (principal components). These eigenvectors represent the directions of motion, while their corresponding eigenvalues indicate the magnitude of these motions. We performed PCA using only backbone atoms employing the gmx covar and gmx anaeig tools from the GROMACS package.

### 2.5 Free energy landscape

In conjunction with PCA, the free energy landscape (FEL) is used to characterize the conformational landscape of the system. The relatively stable conformational states, particularly those near the native state, are represented as local minima on the energy surface [52]. FEL has been widely used to investigate molecular motions and conformational dynamics [53-55]. We selected the first two principal components (PC1 and PC2) as reaction coordinates. The relative free energy difference between two states is given by the following equation:

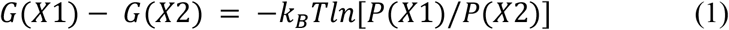

where *X*1and *X*2are the reaction coordinates corresponding to the two PCs, *k*_*B*_ is the Boltzmann constant, T is the temperature, and *P*(*X*1)and *P*(*X*2) represent the probability distributions along these coordinates.

### 2.6 Dynamic cross correlation

Dynamic cross-correlation matrices (DCCMs) are commonly computed by normalizing the covariance matrix of atomic fluctuations. The degree of correlated motion between atoms is quantified by evaluating the correlation between pairs of atoms. The Pearson correlation coefficient [56] is typically used for this purpose and is defined as the following equation:

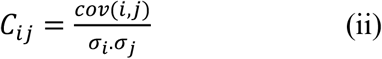

where *C*_*ij*_ is the correlation coefficient between atoms *i* and *j, cov*(*i,j*) represents the covariance of their positions, and σ_*i*_ and σ_*j*_ are the corresponding standard deviations.

### 2.7 Calculation of binding free energy

To calculate the binding affinity of FUS RRM to U1 snRNA, the MM/PBSA (Molecular Mechanics Poisson–Boltzmann Surface Area) was used as implemented in the gmx_MMPBSA tool of GROMACS [57,58]. The binding free energy is given by the following equation:

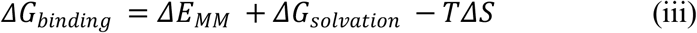

The gas-phase interaction energy Δ*E*_*MM*_ includes internal energy contributed by bond length, bond angle and dihedral angles, as well as van der Waals and electrostatic interactions between the interacting partners. The solvation free energy, Δ*G*_*solvation*_ is decomposed into polar and non-polar components. The polar contribution is calculated using the Poisson–Boltzmann (PB) model, while the non-polar contribution is estimated from the solvent-accessible surface area. The entropic term,−TΔS, is typically obtained from normal-mode analysis of conformational snapshots from MD simulations. The last 500 ns of 1.8 μs simulation was considered for binding energy calculation.

## 3. Results

### 3.1 Structure of FUS RRM bound to U1 snRNA

The 3D structure of FUS RRM bound to U1 snRNA stem loop is shown in Figure 1. The chain A of this protein-RNA complex contains residues 260 to 390 of the 526-residues long FUS. Whereas, chain B is a 28-mer U1 snRNA stem loop. The nIDR region contains 28 residues, and the cIDR region contains 16 residues. The nIDR is part of both RGG1 and RRM domains, whereas the cIDR is included in the RGG2 domain (Figure 1A). The cIDR is bound to the minor groove of snRNA, and the nIDR interacts with the globular part of the protein (Figure 1B). The buried surface area (BSA) contributed by the two IDRs of FUS and that of the RNA is shown in Table 1. The list of interacting atom pairs across the FUS-snRNA interface is tabulated in the supplementary Table S1. Residue stretches 263 to 265 and 283 to 286 in nIDR interact with snRNA.

**Table 1:**
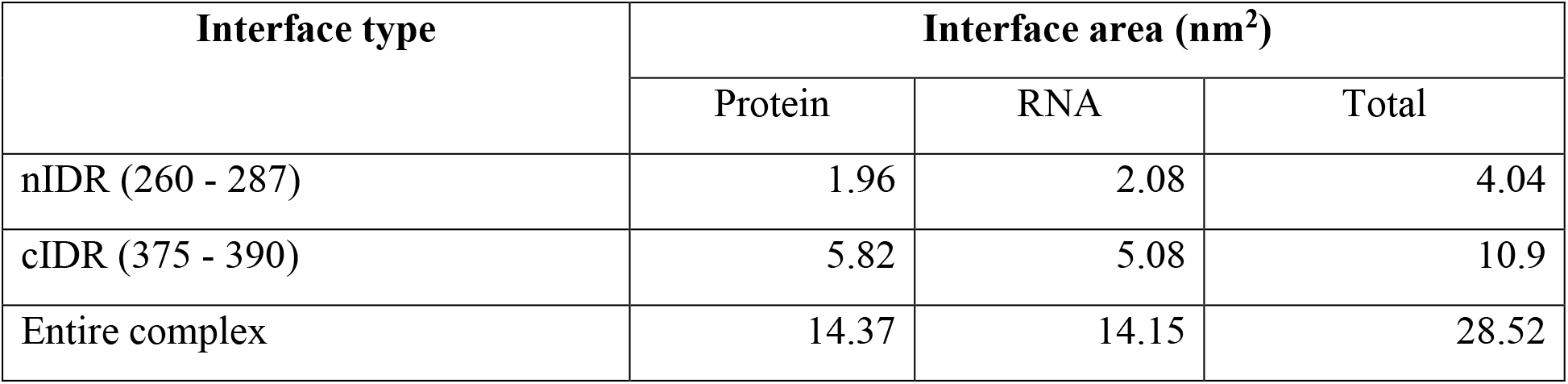
Solvent accessible surface area (SASA) at the protein-RNA interface.

| Interface type | Interface area (nm<sup>2</sup>) |  |  |
| --- | --- | --- | --- |
|  | Protein | RNA | Total |
| nIDR (260 - 287) | 1.96 | 2.08 | 4.04 |
| cIDR (375 - 390) | 5.82 | 5.08 | 10.9 |
| Entire complex | 14.37 | 14.15 | 28.52 |

**Figure 1.**
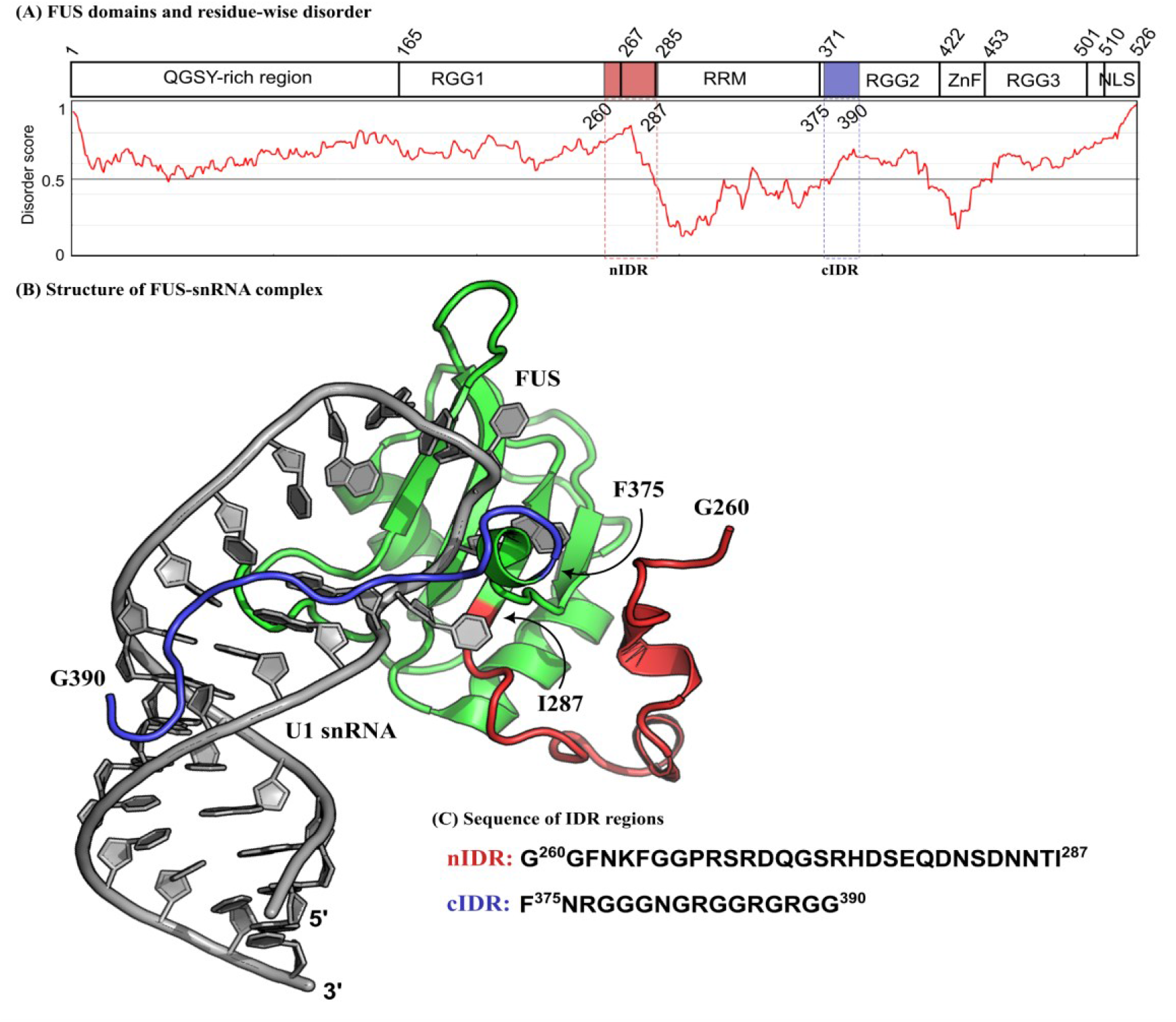
(A) Position of different domains of FUS along its sequence and residue-wise disorder probability score. Two IDRs spanning residues 260 to 287 (nIDR) and 375 to 390 (cIDR), are shown by red and blue rectangles, respectively. (B) Three-dimensional structure of FUS RRM bound to stem loop of U1 snRNA (PDB id: 6SNJ). Protein (chain A) and RNA (chain B) molecules are shown in green and gray cartoons, respectively. The nIDR and the cIDR are shown in red and blue ribbons, respectively. (C) Sequences of nIDR and cIDR.

On the other hand, residue stretches 376 to 377 and 380 to 390 in cIDR interact with the minor groove of the snRNA. The entire protein chain contributes 14.37 nm^2^ to the overall BSA of the complex, of which the two IDRs contribute almost half (7.78 nm^2^). The cIDR (5.82 nm^2^) contributes much more to the overall BSA than the nIDR (1.96 nm^2^). The RNA contributes a total BSA of 14.15 nm^2^, where 7.16 nm^2^ is shared with the two IDRs.

### 3.2 Dynamics of nIDR and cIDR at the RNA interface

Overall structural changes in FUS on binding RNA are monitored by calculating several structural parameters. The backbone RMSD of FUS RRM is compared among the Apo and two replicates of the RNA-bound system. A significant reduction in RMSD is observed in both the complex trajectories (Traj1 and Traj2) compared to Apo FUS (Figure 2A). The time evolution of RMSD is also compared for the protein-RNA complex and the RNA alone in both trajectories (Figure 2B). Traj1 has slightly higher RMSD than Traj2, both for the RNA alone and in the complex. Interestingly, the changes in RMSD in Apo and in RNA-bound forms show different patterns for nIDR and cIDR. RMSD of cIDR of FUS decreases significantly upon binding snRNA, whereas RMSD of nIDR increases slightly upon binding snRNA (Figure 2C). The average RMSD of cIDR decreases from 0.9 nm to 0.3 nm in Traj1 and to 0.5 nm in Traj2. The average RMSD of nIDR in the Apo form increases from 0.6 nm to 0.7 nm (Figure 2D). The residue-level fluctuations (RMSF) clearly show restricted dynamics of cIDR, while the nIDR shows a similar RMSF pattern to that of Apo FUS (Figure 2E). The nucleotides at the loop region of the snRNA (A102 to G108) show higher fluctuation in addition to the nucleotides at the 5′ and 3′ ends (Figure 2F). The radius of gyration of FUS in Traj2 shows similar values to those of the Apo FUS, whereas in Traj1, it is slightly higher. This can be attributed to the extended conformation of cIDR to interact with snRNA (Figure 3A). A comparison of Rg between the FUS-RNA complex and the RNA itself reveals that the snRNA adopts a slightly extended conformation in response to the conformational change of the protein, showing higher Rg in Traj2 (Figure 3B). The overall globular shape of the complex is similar in the two trajectories. Since R_g_ is a measure of compactness of biomolecules, RNA binding reduces the intramolecular compactness of FUS RRM, and facilitates intermolecular interactions.

**Figure 2.**
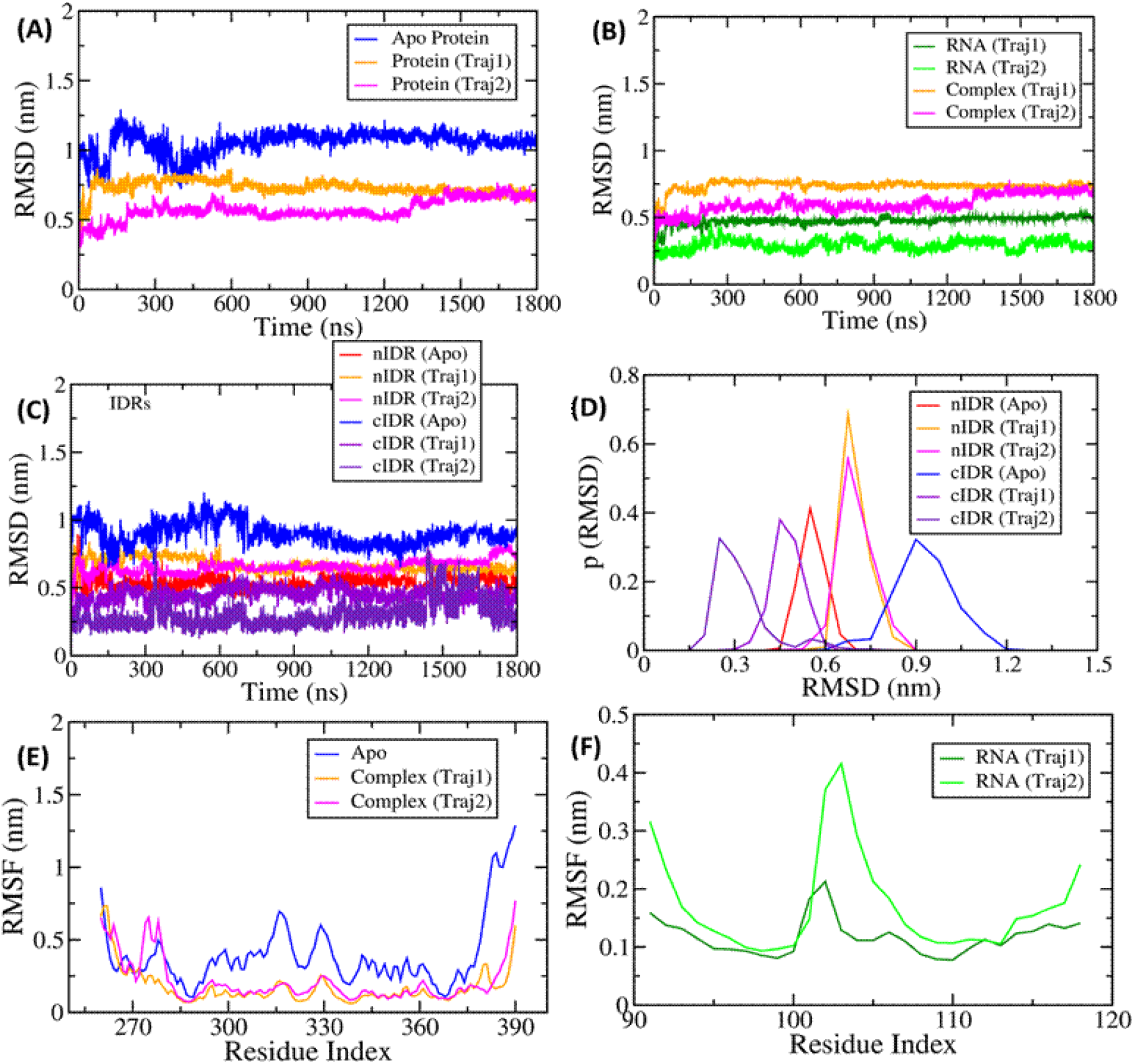
Backbone RMSD and RMSF plots FUS RRM bound to snRNA (PDB id: 6SNJ) (A) RMSD plots of Apo and RNA-bound FUS. (B) RMSD of protein-RNA complexes of two trajectories. (C) RMSD of nIDR and cIDR. (D) Probability density of RMSD for nIDR and cIDR in Apo and two RNA-bound replicates. (E) RMSF plots of Apo and two RNA-bound replicates. (F) RMSF plots of RNA for two RNA-bound systems.

**Figure 3.**
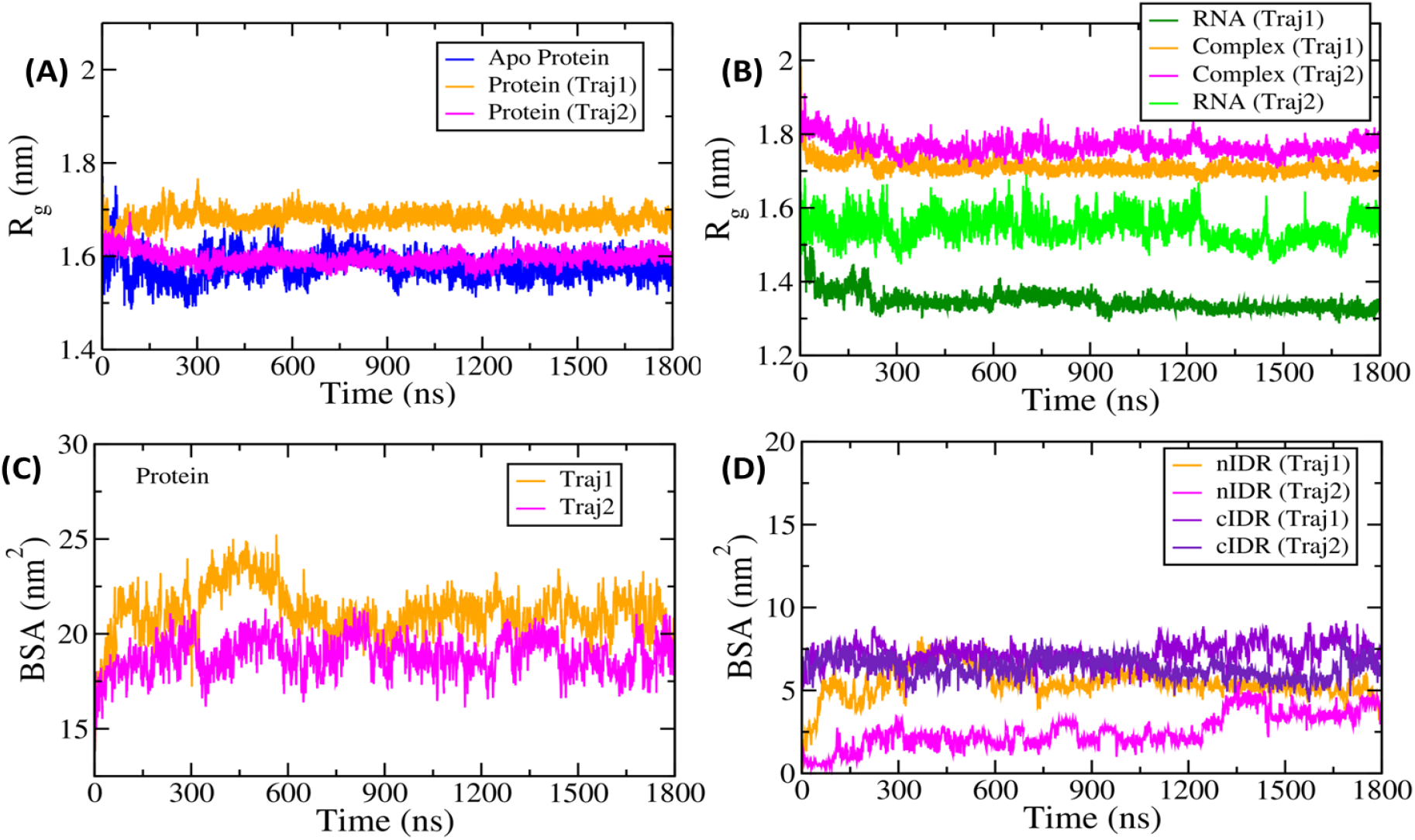
(A) Radius of gyration (R_g_) of Apo and RNA-bound FUS. (B) R_g_ plots for FUS-snRNA complex and snRNA in the two trajectories. (C) Buried surface area (BSA) at the protein-RNA interface of FUS RRM bound to snRNA. (D) BSA contributed by nIDR and cIDR.

Figures 3C and 3D show the BSA contributed by the entire protein chain and by the IDRs, respectively. The BSA of the entire protein chain fluctuates between 15 nm^2^ and 25 nm^2^ (Figure 3C). At the beginning of the simulation, the BSA is within the range of 15 nm^2^ to 20 nm^2^. However, it increases till 600 ns, after which it stabilizes. At 1800 ns, the BSA stays within the range of 17.5 nm^2^ to 20 nm^2^. The BSA of IDRs also shows considerable changes throughout the simulation. The BSA of cIDR shows less fluctuation and remains between 5 nm^2^ and 9 nm^2^ (Figure 3D).

However, BSA of nIDR gradually increases in both trajectories. In Traj1, BSA of nIDR increases to 6 nm^2^ at 300 ns and remains between 5 nm^2^ to 7 nm^2^ in the rest of the simulation. In Traj2, the increase is more gradual and reaches to 5 nm^2^ towards the last 300 ns of the simulation (Figure 3D).

### 3.3 Backbone dihedral angles of IDRs of FUS RRM get rearranged upon RNA binding

The variation in backbone dihedral angles phi (ϕ) and psi (ψ) on a Ramachandran plot indicates the secondary structure of polypeptide chains. The Ramachandran plots of residues of the cIDR and the nIDR are compared between the unbound FUS and in the complex with U1 snRNA (Figure 4). A comparison between the two IDRs in the unbound state reveals that the overall disorderness in the nIDR is lower than the cIDR. Moreover, a significant population of conformations at the beta-sheet and the alpha helical regions is observed in nIDR, indicating the formation of secondary structures during the simulation (Figure 4A(i)). The continuous rearrangement of the backbone dihedral is also persistent in the RNA-bound state, favouring the helical and sheet structures (Figure 4A(ii-iii)). The distribution of the dihedral angle is narrower in cIDR in the unbound state (Figure 4B(i)). Interestingly, conformational space gets even more restricted in cIDR upon binding RNA.The strong interaction of cIDR residues with RNA has restricted the conformational variability, causing a narrower spread in the plot (Figure 4B(ii-iii)). A comparison of these distributions with the backbone dihedral angles from the NMR ensemble (PDB id: 6SNJ) reveals that in addition to the vicinity of the NMR conformations IDRs also sample additional regions of the Ramachandran map.

**Figure 4.**
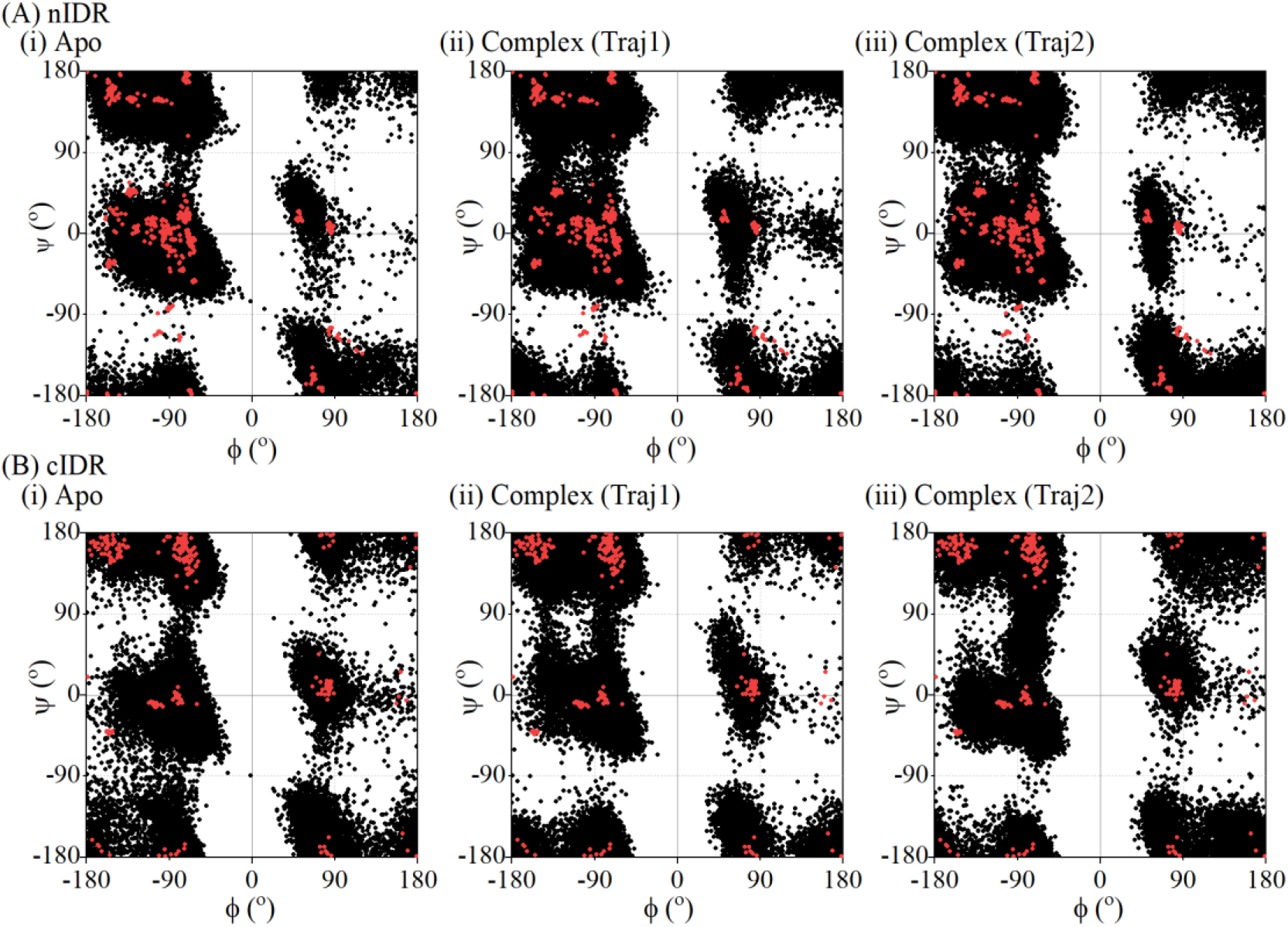
Ramachandran plots of (A) nIDR and (B) cIDR of human FUS in (i) Apo form, (ii) Traj1 and (iii) Traj2. The values corresponding to the NMR ensemble (PDB id: 6SNJ) are plotted in red.

### 3.4 Transition of secondary structures in IDRs upon binding RNA

Figure 5 shows the evolution of secondary structures of FUS RRM in both bound and unbound systems. The terminal IDR stretches (nIDR and cIDR) show an abundance of coils and turns. However, transient α-helices and 3_10_ helices are formed during the simulation, especially at the nIDR and a few residues at the beginning of cIDR (Figure 5A(i)). The formation of helices is induced by inter-helical interactions between residues from nIDR and cIDR (Figure 5A(ii)). In the complex system, these helices are either absent or transiently formed. In Traj1, a shorter nIDR α-helix is formed towards the latter part of the simulation (Figure 5B), which is absent in Traj2 (Figure 5C). The preference for turn and coil over helices at the IDR regions in the presence of RNA may have originated from favourable interactions of RNA with the unstructured conformation of the amino acids. In the ordered regions of the protein, neither the Apo system nor the RNA-bound complex systems show much variation of secondary structures. α-helices and β-sheets in the ordered regions remain unaffected during simulation.

**Figure 5.**
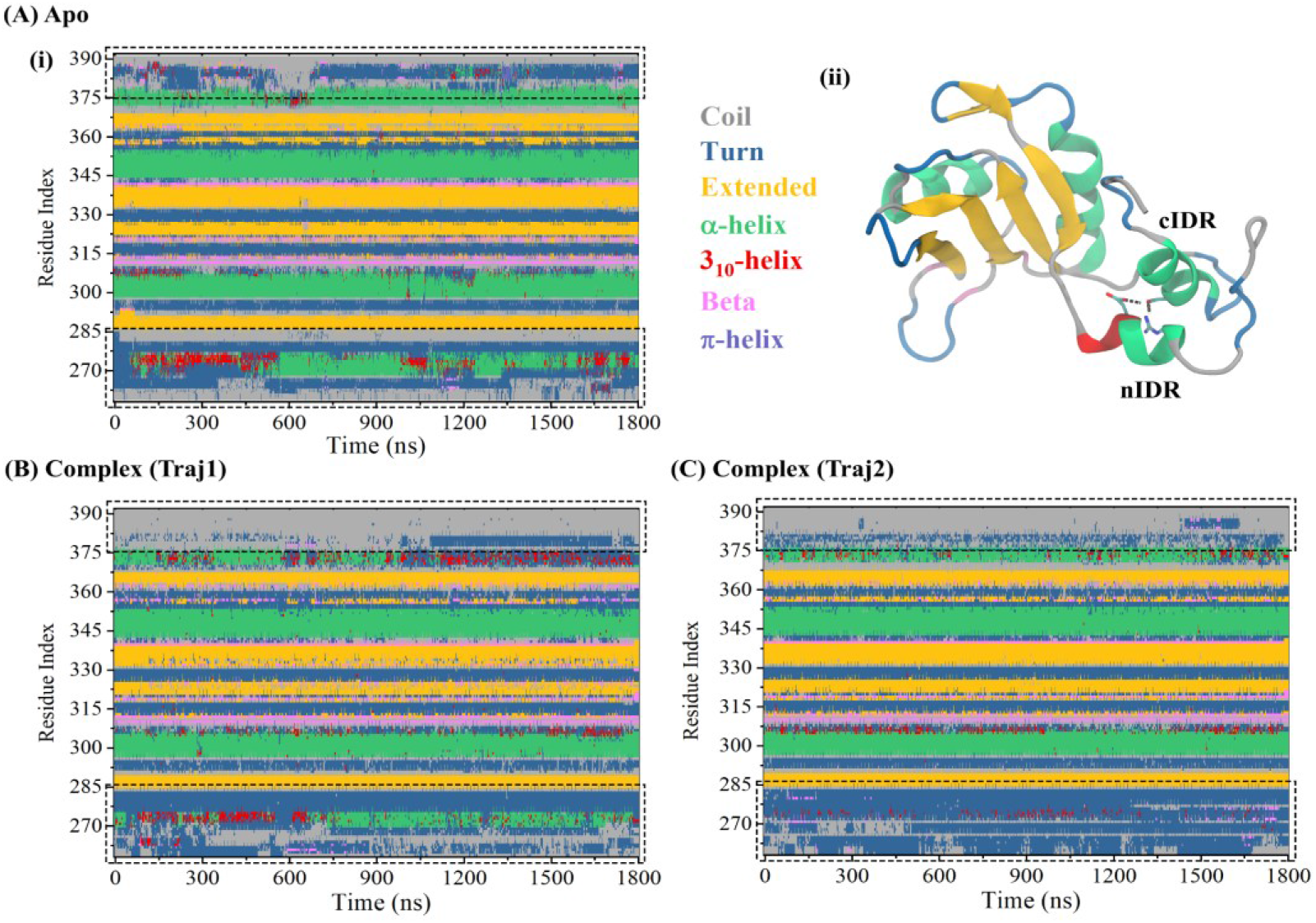
(A) (i) Evolution of secondary structure of residues of FUS RRM in Apo protein. (ii) A representative structure showing interactions between nIDR and cIDR. Evolution of secondary structure in (B) Traj1 and (C) Traj2.

### 3.5 Interactions governing the binding of FUS RRM and U1 snRNA

The binding affinity between FUS and U1 snRNA is measured by calculating the binding energy. In the two trajectories, the affinity values are similar (-77.98 kcal/mol and -70.48 kcal/mol), which highlights the statistical robustness of the results (Table 2).

**Table 2.** Binding energy of FUS RRM bound to snRNA. Energy values are in kJ/mol.

| System | $\Delta E_{\text{vdw}}$ | $\Delta E_{\text{elec}}$ | $\Delta G_{\text{Solv\_nonpolar}}$ | $\Delta G_{\text{Solv\_polar}}$ | $\Delta G_{\text{binding}}$ |
| --- | --- | --- | --- | --- | --- |
| Complex (Traj1) | -234.85±0.42 | -833.20±0.99 | -147.65±0.22 | 990.06±0.98 | -77.98±0.32 |
| Complex (Traj2) | -188.55±0.31 | -770.13±1.40 | -122.28±0.15 | 888.20±1.30 | -70.48±0.31 |

The binding is governed by both electrostatic and van der Waals interactions. To identify important interactions governing the binding, contributions of interface amino acids towards the overall binding affinity are calculated for both trajectories (Figure 6A).

**Figure 6.**
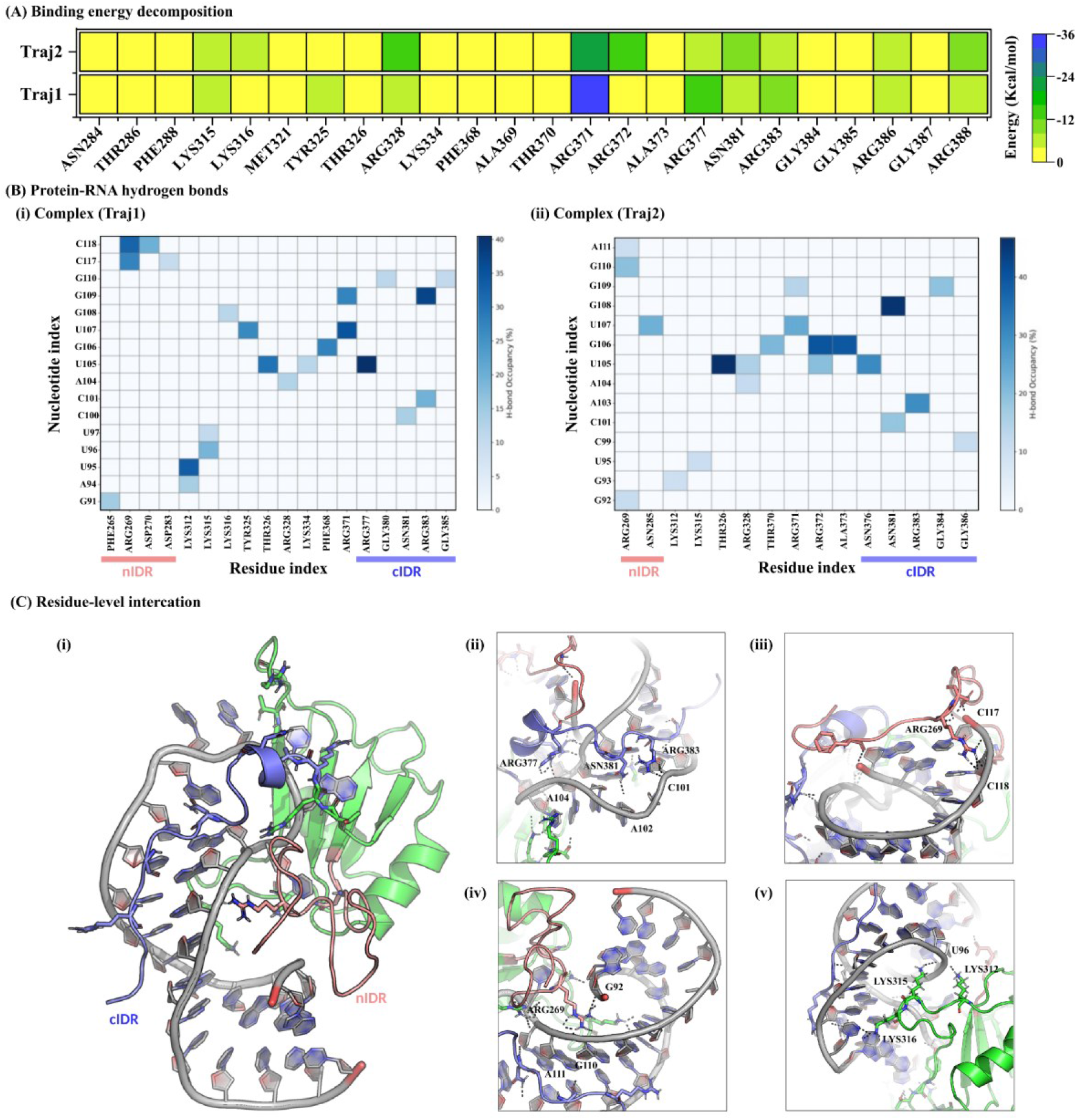
Interactions between residues and nucleotides. (A) Binding energy decomposition for each residue (B) H-bond occupancy heatmaps of the two RNA-bound replicates (i) Traj1 and (ii) Traj2. nIDR and cIDR are indicated by red and blue lines, respectively. (C) Important residue level interactions at the protein-RNA interface. Participating residues are labelled in black. nIDR and cIDR are shown in red and blue cartoons, respectively. The protein and RNA chains are shown in green and gray cartoons, respectively.

Residues of cIDR (ARG371, ARG377, ARG383 and ARG388) contribute significantly to binding along with residues from the RRM domain (LYS315, LYS316, TYR325 and ARG238). The arginine residues in cIDR, contributing significantly to the binding energy, are part of the RGG motifs and are known for their important roles in RNA recognition [59]. The specific protein-RNA interactions stabilizing the FUS-U1 snRNA complex are identified as H-bond interaction between the interface residues and nucleotides (Figure 6B). Figure 6B(i) and 6B(ii) show the heatmaps of the occupancy of H-bonds in Traj1 and Traj2, respectively. The occupancy of the H-bond is expressed as a percentage of time the bond exists in the total period of simulation. The occupancy values of all H-bonds in Traj1 and Traj2 are listed in Supplementary Table S2. In both the trajectories, the nIDR mainly forms H-bonds with G91 to C99 at the 5′ end and with A113 to C118 at the 3′ end of the snRNA. On the other hand, the cIDR interacts with the stem-loop region (C100 to A112) of the U1 snRNA. A careful inspection of the trajectories reveals that the cIDR spans along the minor groove of the U1 snRNA stabilized by interaction of polar side chains with the phosphate and sugar backbone of the RNA. On the other hand, nIDR interacts with the terminals of the snRNA with several transient protein-RNA interactions (Figure 6C(i)). ARG 377, ARG381 and ARG383 from cIDR are involved in H-bonds with the backbone atoms of A104, A102 and C101, respectively (Figure 6C(ii)). ARG269 of nIDR plays a crucial role in coordinating U1 snRNA, however the point of interaction differs in the two trajectories. In Traj1, RG269 interacts with C117 and C118 (Figure 6C(iii)), and in Traj2, it bridges G92 and G110 (Figure 6C(iv)). Other than nIDR and cIDR, LYS312 and LYS315 are also involved in interaction with RNA-backbone for a significant simulation time (Figure 6C(v)).

### 3.6 PCA and FEL of Apo and RNA-bound FUS

To explore the dominant modes of motion during the simulation, PCA was performed for Apo and RNA-bound FUS (Figure 7). The contribution of the resulting principal components is shown in Figure S1. The first two principal components (PC1 and PC2) constitute more than 50% of the dynamic behaviour in all systems. The ensemble of conformations is projected on PC1 and PC2 to understand the conformational landscape. Figure 7A corresponds to the Apo protein, where the distribution is broad and continuous. The points are spread over a wide region, and the major clusters are not well-separated into distinct clusters, suggesting that FUS samples a large conformational space and remains quite flexible in the absence of snRNA.

**Figure 7.**
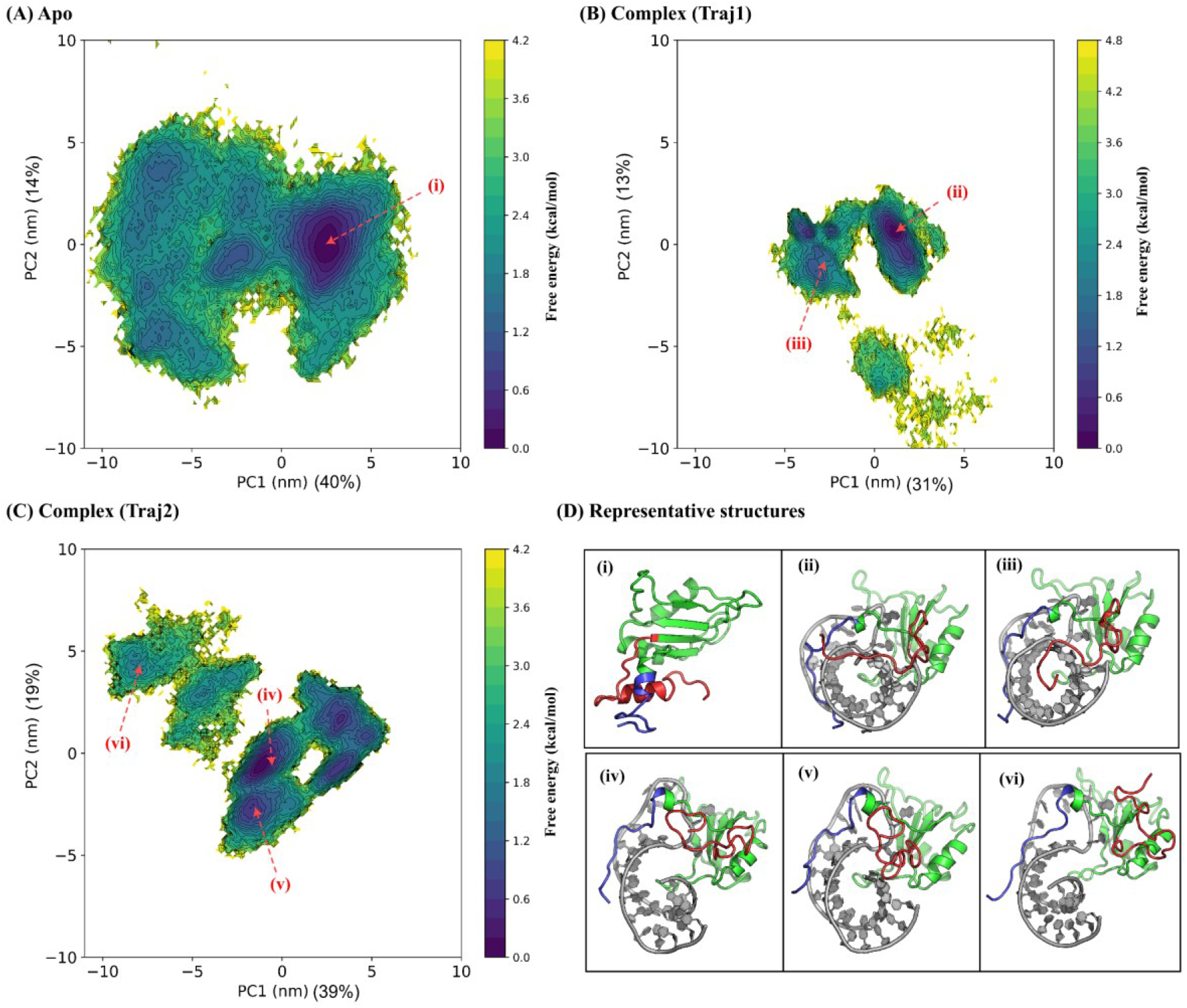
PCA of the FUS RRM bound to snRNA. (A) Apo protein showing broad and continuous conformational sampling. (B and C) Two independent simulations of the protein-RNA complex showing multiple basins, indicating restricted and structured dynamics upon binding RNA. D (i) to (vi) show structures at different minima marked by arrows in A-C.

It has one dense central region or global minimum (shown in dark blue (i)), representing the most populated conformations, while the outer scattered regions represent less frequent states. A representative conformation from the minimum depicts tertiary interactions between cIDR and nIDR (Fig. 7D(i)). The two independent replicates of the protein-RNA complex show a completely different picture (Figure 7B-C). Instead of continuous distributions, the conformations are grouped into multiple distinct clusters of small conformational subspace, suggesting relatively restricted motion and stabilization of few selective conformations. These clusters suggest that the protein has fewer preferred conformational states when binds RNA. The presence of separated basins indicates transitions between stable states, which are more defined compared to the Apo. Among the two replicates, Traj1 shows two local minima separated along PC1, along with some minor clusters at a lower PC2 range, indicating large conformational transitions (Figure 7B). The representative conformations from these two minima are shown in Figure 7D(ii) and (iii). In both cases, the cIDR interacts with the minor groove of the snRNA, while the nIDR either spans over the 5′ and 3′ ends Figure 7D(ii) or anchors into the major groove Figure 7D(iii). PC1-PC2 distribution in Traj2 has four major clusters and a minor, less populated region. The representative structures corresponding to two local minima have different nIDR-RNA interactions. One of them is more interacting with the RRM region, with a few interactions with the stem-loop part Figure 7D(iv). In the other minimum, nIDR is strongly involved in interactions with the major groove Figure 7D(v). The third isolated cluster represents a relatively less compact complex with nIDR projected away from the snRNA and involved in intra-chain interactions (vi). Overall, the RNA binding reduces the conformational freedom of the FUS and organizes its motion into specific functional states.

### 3.7 Dynamic cross correlation

Figure 8 presents the correlation maps between C_α_ atoms for the Apo and the complex systems. Strong positive correlations are visible for unbound FUS, which represent coordinated motions between specific residue pairs in the free form (shown as red rectangles in Figure 8A). Such correlations reflect dynamic communication within the structure. In case of the RNA-bound state (Traj1 and Traj2), such correlations are reduced (Figure 8B and 8C), as indicated by the disappearing regions inside the red rectangles. In the Apo system, residues 266 to 285 in the nIDR show cross-correlation with residues 300 to 320. This correlation is absent in the RNA-bound complex. Similarly, correlation between residues 300 to 320 with residues 360 to 380, which overlaps with the cIDR, is seen in the Apo system. However, this correlation disappears in the RNA-bound complex. This indicates that RNA binding disrupts the coordinated motions between disordered residues, impairing the dynamic coupling present in the unbound state. The loss of these correlations suggests that RNA interaction reshapes the internal communication network of the protein. This suggests RNA binding does not simply add new contacts but actively interferes with pre-existing residue correlations.

**Figure 8.**
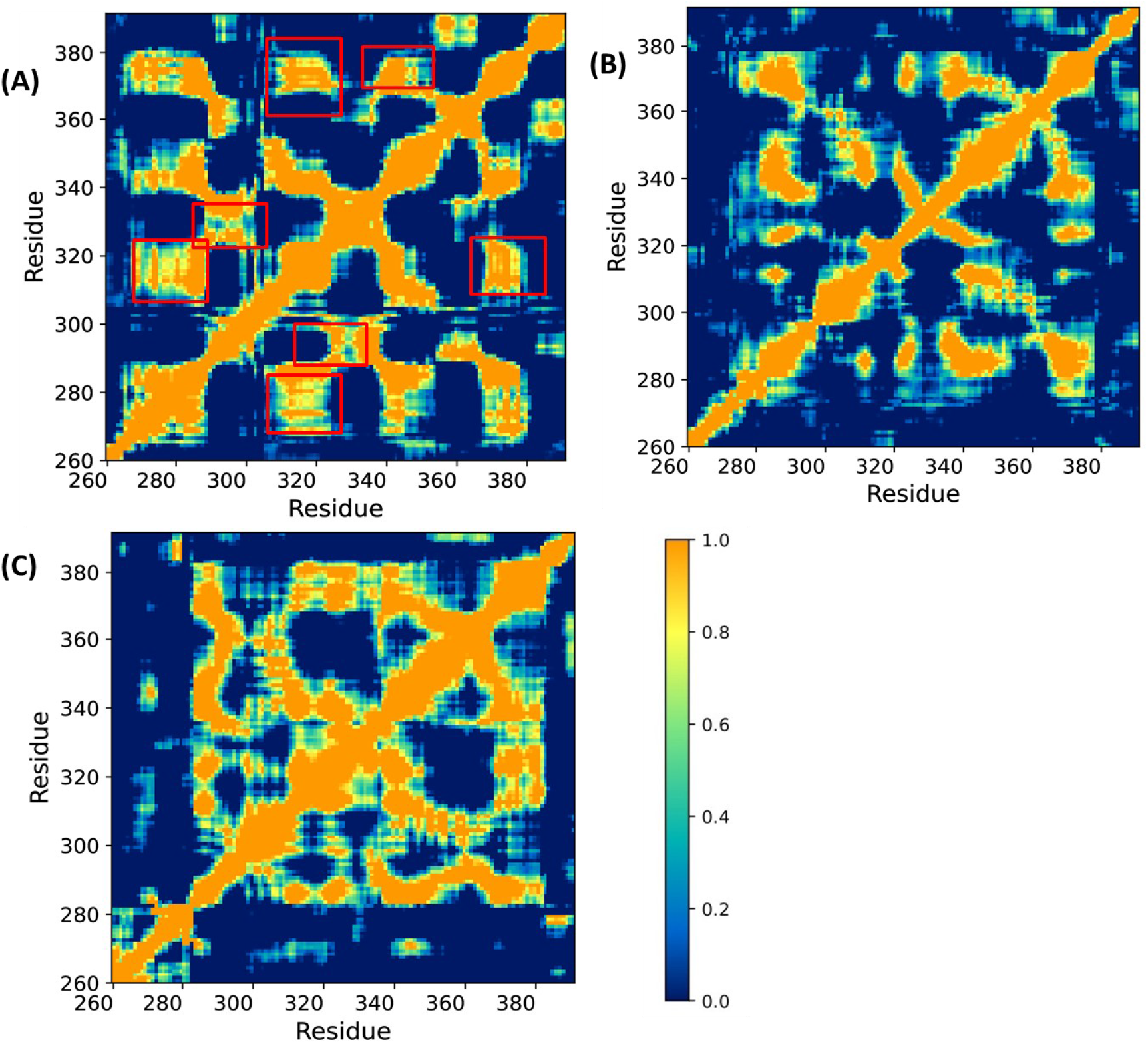
DCCM plots of C_a_ atoms of FUS RRM bound to snRNA (PDB id: 6SNJ) of (A) Apo system and (B) complex (Traj1) and (C) complex (Traj2). Regions of strong cross-correlation are demarcated by red rectangles which are missing upon binding RNA.

To understand the direction of movement of the components during the simulation, the porcupine plots are generated by considering the two extreme states based on PC1 using gmx anaeig (Figure 9). The arrows show the direction of the motion, and the size of the arrows represents how much that part of the protein moved from its original position. For the Apo protein (Figure 9A), the arrows are spread over almost the whole structure. Some arrows point in different directions, which means different parts of the protein move independently, but have higher correlations. Some regions have large displacements, especially in the two IDRs and loop regions. This is expected since these regions are flexible. The cIDR (GLY385, ARG386, GLY387, ARG388, GLY389 and GLY390) has comparatively higher displacement. In the case of RNA-bound complexes (Figure 9B and 9C), the pattern changes. The arrows are no longer spread everywhere but are mostly concentrated in specific regions (IDRs and loops). This means that RNA binding restricts the motion of the protein in some regions and stabilizes the complex.

**Figure 9.**
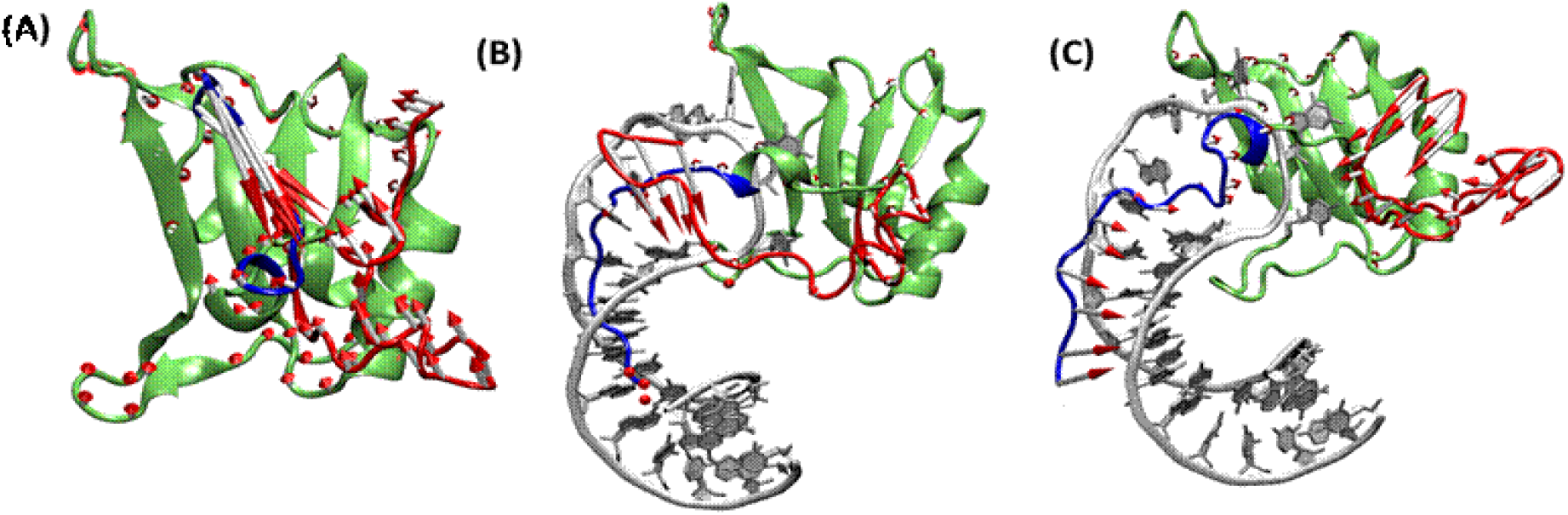
Porcupine plots indicating dominant motions along the first principal component: (A) Apo protein showing widespread and random motions. (B and C) Two replicates of protein-RNA complex displaying more localized and directional movements. The protein is shown in green color in cartoon representation, while the RNA is shown in secondary structure representation in grey. The nIDR and the cIDR are shown in red and blue, respectively.

## 4. Discussion

### RNA-bound IDRs of FUS show distinct and salient dynamics compared to unbound FUS

Our results show that the IDRs in FUS behave in a distinct manner when interacting with the partner snRNA. This difference is reflected in various interactions and dynamic behaviour of the residues that constitute disordered stretches. IDRs of FUS show frequent change in interaction surface with U1 snRNA, as indicated by higher fluctuations in BSA during simulation. This is also supported by a plethora of H-bond interactions between the residues in IDRs and the nucleotides in snRNA. A comparison of backbone dihedral angles and secondary structures of the IDR residues in Apo and RNA-bound states depicts that some nIDR residues transiently adopt α-helices and 3_10_ helices, whereas residues in cIDR remain as turns or coils. Binding with the partner U1 snRNA limits the adoption of helical secondary structures in IDRs. In the Apo state, the overall disorderness in the cIDR is significantly higher than in the nIDR, as supported by the spread of dihedral angles. RNA binding causes continuous rearrangement of dihedral angles of IDR residues. In nIDR, RNA binding slightly favours the helical region, whereas the distribution of dihedral angles of cIDR gets even more stringent upon binding RNA due to the association of the C-terminal region in the RNA groove. Interestingly, the population of ‘turn’ is significantly reduced in cIDR upon binding RNA. This can be attributed to the inter-residue interaction in the Apo state, which is replaced by strong interactions with RNA minor groove in the complex. The absence of large-scale disorder-to-order transitions in RGG1 (part of nIDR) is observed in a combined NMR and EPR investigation by Bonucci et al., [60]. Structural deviation of residues in IDRs also decreases upon RNA binding due to its stabilizing effect as supported by the RMSD and RMSF plots. Despite cIDR being smaller in size than nIDR, the BSA between cIDR and RNA is higher than that between nIDR and RNA (Table 1). During simulation, BSA of nIDR gradually increases, highlighting increased interaction between nIDR and RNA. The interaction is also reflected in the lifetime of H-bonds at the protein-RNA interface (Figure 6B). Decomposition of binding energy as well as analysis of intra-chain H-bonds also reveal that IDRs have much favourable binding energy than the ordered stretches. Residues in cIDR interact more with the RNA minor groove, while certain nIDR residues tend to recognize terminal nucleotides.

### RNA binding alters the correlated movements of IDRs

The correlation maps (Fig. 8) and porcupine plots (Fig. 9) together show a clear picture of how RNA binding changes the dynamic behaviour of the FUS. In the Apo form, the protein shows strong coordinated movements between different regions. These are seen as clear correlated zones between residues, suggesting that these segments of the protein are dynamically correlated. In particular, residues in the nIDR region move in a coordinated way with the residues in the loop region. Similarly, another connection exists between residues 300 to 320 and the residues in the cIDR (Figure 8A). Upon binding RNA, these coordinated movements are weakened and these correlations disappear. This indicates that RNA binding disrupts the internal communication between N- and C-terminal regions. Instead of moving together, different parts of the protein show restricted motion. The porcupine plots (Figure 9) further support this observation by showing how the actual movements of the protein parts change. In the Apo protein, motion is widespread with different regions moving in different directions but in a coordinated manner within a segment. This reflects a flexible system where multiple parts contribute to the overall dynamics of FUS. The non-specificity and multivalent nature of IDR-RNA interaction is also reported in a computational investigation of concerted binding of RGG and ZnF domains of FUS [61]. The largest movements are seen in the disordered regions, especially in cIDR, which appears to be the most mobile segment. In the complex systems, the motion becomes more localized. Instead of the whole protein moving freely, only certain regions, mainly loops and disordered segments show significant displacement. Even within these regions, the movement is more controlled and less scattered. However, some parts of the disordered regions nIDR and cIDR retain their flexibility.

### RNA binding narrows the conformational landscape of the protein

From the PCA results we observe that the overall behavior of the protein changes with RNA binding (Fig. 7). In the Apo state, the protein explores a wide range of conformations, which suggests the structure is not confined to a few conformational states but distributed across different conformational ensembles. In contrast, the RNA-bound system shows a more organized motion. In this case, the protein tends to occupy a limited but multiple number of clusters. It means the system interconverts between a few stable states rather than sampling continuously. The RNA introduces structural constraints and guides the protein to adopt specific conformational states. Similar RNA binding-induced restriction and redistribution of protein conformational dynamics have been reported previously in NMR and simulation studies of protein–RNA complexes [62,63]. The cumulative variance also supports this observation from a different perspective. In the Apo system, the motion is distributed across many components, and no single or small group of motions dominates the dynamics. On the other hand, in the RNA-bound systems, a larger fraction of the total motion is captured within fewer components, and the dynamics are dominated by a limited number of collective movements. Such behaviour is often associated with more dynamically constrained systems, where the motion is not random, rather follows certain preferred directions.

## 5. Conclusion

In this study, we investigate the dynamics of the intrinsically disordered regions (IDRs) flanking the RRM domain of human FUS in both Apo and U1 snRNA-bound states using molecular dynamics simulations. Comparison of hydrogen-bonded interactions, buried surface area, secondary structure transitions and correlated motions reveals that RNA binding stabilizes the IDRs and restricts their conformational dynamics by reorganizing intra-protein contacts. The N-terminal and the C-terminal IDRs exhibit distinct modes of RNA recognition. While nIDR interacts primarily with terminal nucleotides (from G91 to C99 and from A113 to C118) of the stem-loop RNA, the cIDR preferentially associates with the minor groove of the central double-stranded region (C100 to A112) through interactions with the RNA backbone. Together, these findings provide molecular-level insights into the dynamics and functionally distinct roles of two flanking disordered regions around the RRM domain of FUS in recognizing stem-loop snRNA.

## Acknowledgement

S.M. and F.K.M. acknowledge the fellowship and infrastructure provided by Indian Institute of Technology Kharagpur. RPB acknowledges IIT Kharagpur for infrastructure support. AM and RPB acknowledge DBT, Govt. of India for computational facility funded by the BIC programme (Grant no. BT/PR40175/BTIS/137/41/2022). All the authors are thankful to DBT for providing the computational facility and the National Supercomputing Mission (NSM) for providing computing resources of ‘PARAM Shakti’ at IIT Kharagpur, supported by the Department of Science and Technology (DST), Government of India.

## Author contributions

S.M. conceptualized the work, formal analysis and wrote the initial draft, review and editing. F.K.M. performed the simulations, formal analysis, writing, review and editing. A.M. did analysis, visualization, and writing, review and editing of the manuscript. R.P.B. conceptualized the work, analyzed data, reviewed and edited the manuscript, overall supervised the project, and arranged funding and resources.

