## Supplementary information for "Deciphering the role of intrinsically disordered regions of FUS in recognizing U1 snRNA"

**for**

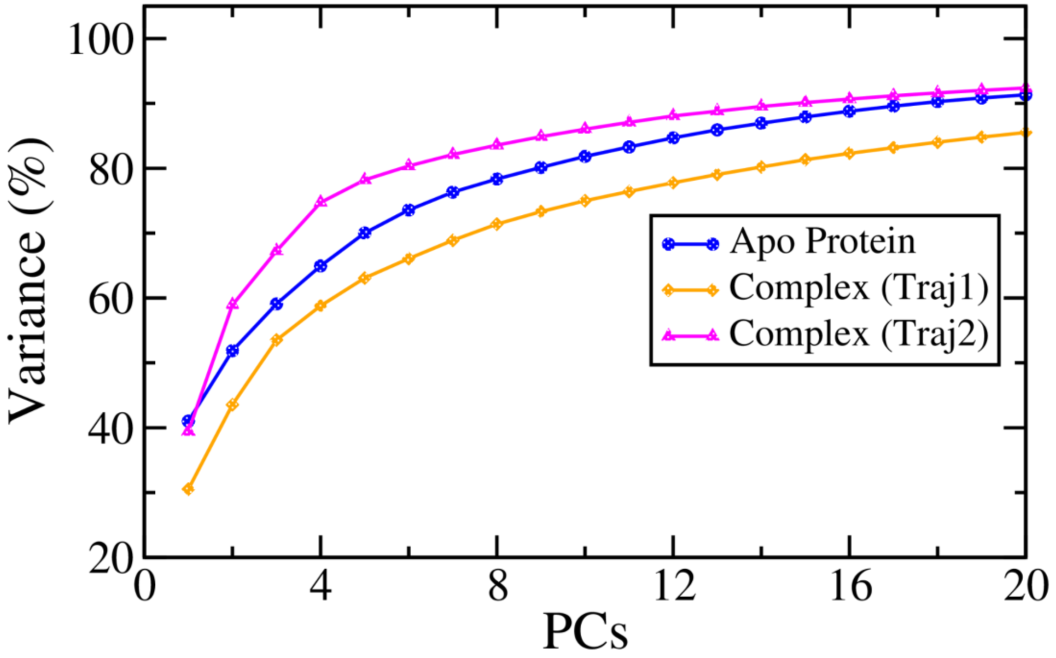


Supplementary Figure S1: Cumulative variance of principal components (PCs) for the apo and two RNA-bound replicates (Traj1 and Traj2).

Supplementary Table S1: Interacting atom pairs between the FUS and the U1 snRNA within a cutoff of distance of 4.5 Å.

| IDR name | Protein atom name | Residue name | Residue number | RNA atom name | Nucleotide | Nucleotide number | Distance |
|---|---|---|---|---|---|---|---|
| nIDR | CB | ASN | 263 | N2 | G | 106 | 4.3 |
| nIDR | CG | ASN | 263 | N1 | G | 106 | 3.7 |
| nIDR | CG | ASN | 263 | C2 | G | 106 | 3.9 |
| nIDR | CG | ASN | 263 | N2 | G | 106 | 3.3 |
| nIDR | OD1 | ASN | 263 | C6 | G | 106 | 3.8 |
| nIDR | OD1 | ASN | 263 | O6 | G | 106 | 3.9 |
| nIDR | OD1 | ASN | 263 | N1 | G | 106 | 2.8 |
| nIDR | OD1 | ASN | 263 | C2 | G | 106 | 3.4 |
| nIDR | OD1 | ASN | 263 | N2 | G | 106 | 3.1 |
| nIDR | ND2 | ASN | 263 | N1 | G | 106 | 4.3 |
| nIDR | ND2 | ASN | 263 | C2 | G | 106 | 4.3 |
| nIDR | ND2 | ASN | 263 | N2 | G | 106 | 3.2 |
| nIDR | D21 | ASN | 263 | N1 | G | 106 | 4.2 |
| nIDR | D21 | ASN | 263 | C2 | G | 106 | 4.1 |
| nIDR | D21 | ASN | 263 | N2 | G | 106 | 3 |
| nIDR | D22 | ASN | 263 | N2 | G | 106 | 3.9 |
| nIDR | NZ | LYS | 264 | C5 | G | 106 | 4.4 |
| nIDR | NZ | LYS | 264 | C6 | G | 106 | 3.7 |
| nIDR | NZ | LYS | 264 | O6 | G | 106 | 3.4 |
| nIDR | NZ | LYS | 264 | N1 | G | 106 | 4.3 |
| nIDR | CE2 | PHE | 265 | C2 | G | 106 | 4.3 |
| nIDR | CE2 | PHE | 265 | N2 | G | 106 | 3.6 |
| nIDR | CZ | PHE | 265 | N2 | G | 106 | 3.7 |

| IDR name | Protein atom name | Residue name | Residue number | RNA atom name | Nucleotide | Nucleotide number | Distance |
|---|---|---|---|---|---|---|---|
| nIDR | C | ASP | 283 | N3 | U | 107 | 3.8 |
| nIDR | C | ASP | 283 | C4 | U | 107 | 4.4 |
| nIDR | C | ASP | 283 | O4 | U | 107 | 4.1 |
| nIDR | O | ASP | 283 | C2 | U | 107 | 3.7 |
| nIDR | O | ASP | 283 | O2 | U | 107 | 3.9 |
| nIDR | O | ASP | 283 | N3 | U | 107 | 2.7 |
| nIDR | O | ASP | 283 | C4 | U | 107 | 3.3 |
| nIDR | O | ASP | 283 | O4 | U | 107 | 3.1 |
| nIDR | N | ASN | 284 | N3 | U | 107 | 4.4 |
| nIDR | CA | ASN | 284 | N3 | U | 107 | 4.2 |
| nIDR | CA | ASN | 284 | C4 | U | 107 | 4.3 |
| nIDR | CA | ASN | 284 | O4 | U | 107 | 3.9 |
| nIDR | CB | ASN | 284 | O4 | U | 107 | 4.3 |
| nIDR | CG | ASN | 284 | C4 | U | 107 | 4.4 |
| nIDR | CG | ASN | 284 | O4 | U | 107 | 3.4 |
| nIDR | OD1 | ASN | 284 | O4 | U | 107 | 3.6 |
| nIDR | ND2 | ASN | 284 | C4 | U | 107 | 4 |
| nIDR | ND2 | ASN | 284 | O4 | U | 107 | 3 |
| nIDR | D21 | ASN | 284 | C4 | U | 107 | 3.9 |
| nIDR | D21 | ASN | 284 | O4 | U | 107 | 2.8 |
| nIDR | D22 | ASN | 284 | C4 | U | 107 | 4.2 |
| nIDR | D22 | ASN | 284 | O4 | U | 107 | 3.4 |
| nIDR | OD1 | ASN | 285 | C2 | U | 107 | 4.1 |
| nIDR | OD1 | ASN | 285 | O2 | U | 107 | 3.6 |
| nIDR | OD1 | ASN | 285 | N3 | U | 107 | 4.4 |
| nIDR | D21 | ASN | 285 | O2 | U | 107 | 4.2 |

| IDR name | Protein atom name | Residue name | Residue number | RNA atom name | Nucleotide | Nucleotide number | Distance |
|---|---|---|---|---|---|---|---|
| nIDR | CB | THR | 286 | O4' | U | 107 | 4.1 |
| nIDR | CB | THR | 286 | N1 | U | 107 | 4.1 |
| nIDR | CB | THR | 286 | C5 | U | 107 | 3.9 |
| nIDR | CB | THR | 286 | C6 | U | 107 | 3.7 |
| nIDR | OG1 | THR | 286 | O4' | U | 107 | 3.2 |
| nIDR | OG1 | THR | 286 | C1' | U | 107 | 3.6 |
| nIDR | OG1 | THR | 286 | N1 | U | 107 | 3.2 |
| nIDR | OG1 | THR | 286 | C2 | U | 107 | 3.7 |
| nIDR | OG1 | THR | 286 | O2 | U | 107 | 4.3 |
| nIDR | OG1 | THR | 286 | N3 | U | 107 | 4.2 |
| nIDR | OG1 | THR | 286 | C4 | U | 107 | 4.3 |
| nIDR | OG1 | THR | 286 | C5 | U | 107 | 3.8 |
| nIDR | OG1 | THR | 286 | C6 | U | 107 | 3.3 |
| nIDR | CG2 | THR | 286 | O4' | U | 107 | 4.2 |
| nIDR | CG2 | THR | 286 | C6 | U | 107 | 4.2 |
| nIDR | G21 | THR | 286 | O4' | U | 107 | 4.3 |
| nIDR | G22 | THR | 286 | O3' | G | 106 | 4.2 |
| nIDR | G22 | THR | 286 | O2' | G | 106 | 4 |
| nIDR | G22 | THR | 286 | O2' | G | 106 | 4.1 |
| nIDR | G22 | THR | 286 | C5' | U | 107 | 3.9 |
| nIDR | G22 | THR | 286 | C4' | U | 107 | 4.3 |
| nIDR | G22 | THR | 286 | O4' | U | 107 | 3.5 |
| nIDR | G22 | THR | 286 | N1 | U | 107 | 4.3 |
| nIDR | G22 | THR | 286 | C5 | U | 107 | 4.2 |
| nIDR | G22 | THR | 286 | C6 | U | 107 | 3.6 |
| cIDR | CG | ASN | 376 | OP1 | U | 105 | 3.9 |

| IDR name | Protein atom name | Residue name | Residue number | RNA atom name | Nucleotide | Nucleotide number | Distance |
|---|---|---|---|---|---|---|---|
| cIDR | OD1 | ASN | 376 | P | U | 105 | 4.2 |
| cIDR | OD1 | ASN | 376 | OP1 | U | 105 | 3.7 |
| cIDR | OD1 | ASN | 376 | OP2 | U | 105 | 3.7 |
| cIDR | ND2 | ASN | 376 | OP1 | U | 105 | 3.3 |
| cIDR | D21 | ASN | 376 | P | U | 105 | 3.6 |
| cIDR | D21 | ASN | 376 | OP1 | U | 105 | 2.3 |
| cIDR | D21 | ASN | 376 | OP2 | U | 105 | 4 |
| cIDR | D21 | ASN | 376 | O5' | U | 105 | 4.4 |
| cIDR | D22 | ASN | 376 | OP1 | U | 105 | 4 |
| cIDR | CG | ARG | 377 | O3' | A | 104 | 4.4 |
| cIDR | CG | ARG | 377 | P | U | 105 | 4 |
| cIDR | CG | ARG | 377 | OP1 | U | 105 | 3.5 |
| cIDR | CG | ARG | 377 | OP2 | U | 105 | 3.8 |
| cIDR | CD | ARG | 377 | P | U | 105 | 4.4 |
| cIDR | CD | ARG | 377 | OP1 | U | 105 | 4.4 |
| cIDR | CD | ARG | 377 | OP2 | U | 105 | 3.7 |
| cIDR | CZ | ARG | 377 | P | A | 104 | 4.4 |
| cIDR | CZ | ARG | 377 | OP1 | A | 104 | 3.6 |
| cIDR | CZ | ARG | 377 | O5' | A | 104 | 4.1 |
| cIDR | CZ | ARG | 377 | C5' | A | 104 | 4 |
| cIDR | CZ | ARG | 377 | C3' | A | 104 | 4.4 |
| cIDR | CZ | ARG | 377 | OP2 | U | 105 | 4.3 |
| cIDR | NH1 | ARG | 377 | P | A | 104 | 3.8 |
| cIDR | NH1 | ARG | 377 | OP1 | A | 104 | 3.5 |
| cIDR | NH1 | ARG | 377 | OP2 | A | 104 | 4.2 |
| cIDR | NH1 | ARG | 377 | O5' | A | 104 | 3.3 |

| IDR name | Protein atom name | Residue name | Residue number | RNA atom name | Nucleotide | Nucleotide number | Distance |
|---|---|---|---|---|---|---|---|
| cIDR | NH1 | ARG | 377 | C5' | A | 104 | 3.5 |
| cIDR | NH1 | ARG | 377 | C4' | A | 104 | 4 |
| cIDR | NH1 | ARG | 377 | C3' | A | 104 | 3.4 |
| cIDR | NH1 | ARG | 377 | O3' | A | 104 | 4 |
| cIDR | NH1 | ARG | 377 | P | U | 105 | 4.2 |
| cIDR | NH1 | ARG | 377 | OP2 | U | 105 | 3.3 |
| cIDR | NH2 | ARG | 377 | P | A | 104 | 4 |
| cIDR | NH2 | ARG | 377 | OP1 | A | 104 | 2.8 |
| cIDR | NH2 | ARG | 377 | O5' | A | 104 | 4.1 |
| cIDR | NH2 | ARG | 377 | C5' | A | 104 | 4 |
| cIDR | CA | GLY | 380 | O2' | G | 108 | 4.2 |
| cIDR | CA | GLY | 380 | O2' | G | 108 | 4.3 |
| cIDR | N | ASN | 381 | O2' | G | 108 | 4.3 |
| cIDR | N | ASN | 381 | O2' | G | 108 | 4.4 |
| cIDR | O | ASN | 381 | O2' | G | 108 | 4.2 |
| cIDR | O | ASN | 381 | N3 | G | 108 | 4.4 |
| cIDR | O | ASN | 381 | O4' | G | 109 | 4.2 |
| cIDR | O | ASN | 381 | C1' | G | 109 | 4.4 |
| cIDR | N | GLY | 382 | O2' | G | 109 | 4.1 |
| cIDR | CA | GLY | 382 | O4' | G | 109 | 4.4 |
| cIDR | CA | GLY | 382 | C2' | G | 109 | 4.2 |
| cIDR | CA | GLY | 382 | O2' | G | 109 | 3.2 |
| cIDR | CA | GLY | 382 | C1' | G | 109 | 4 |
| cIDR | CA | GLY | 382 | O2' | G | 109 | 3.9 |
| cIDR | C | GLY | 382 | O2' | G | 109 | 4.4 |
| cIDR | N | ARG | 383 | N2 | G | 108 | 4.4 |

| IDR name | Protein atom name | Residue name | Residue number | RNA atom name | Nucleotide | Nucleotide number | Distance |
|---|---|---|---|---|---|---|---|
| cIDR | N | ARG | 383 | N3 | G | 109 | 4.2 |
| cIDR | CA | ARG | 383 | N2 | G | 109 | 4.4 |
| cIDR | C | ARG | 383 | N2 | G | 109 | 3.8 |
| cIDR | O | ARG | 383 | C2 | G | 109 | 4.1 |
| cIDR | O | ARG | 383 | N2 | G | 109 | 3.2 |
| cIDR | O | ARG | 383 | N3 | G | 109 | 4.1 |
| cIDR | CB | ARG | 383 | C2 | C | 100 | 4.1 |
| cIDR | CB | ARG | 383 | O2 | C | 100 | 3 |
| cIDR | CB | ARG | 383 | N2 | G | 108 | 3.4 |
| cIDR | CB | ARG | 383 | N2 | G | 109 | 4.2 |
| cIDR | CG | ARG | 383 | O2 | C | 100 | 3.3 |
| cIDR | CG | ARG | 383 | N3 | C | 101 | 4 |
| cIDR | CG | ARG | 383 | C2 | G | 108 | 4.4 |
| cIDR | CG | ARG | 383 | N2 | G | 108 | 3.1 |
| cIDR | CD | ARG | 383 | O2 | C | 100 | 4.4 |
| cIDR | CD | ARG | 383 | C2 | C | 101 | 4.4 |
| cIDR | CD | ARG | 383 | O2 | C | 101 | 4 |
| cIDR | CD | ARG | 383 | N3 | C | 101 | 4.3 |
| cIDR | NE | ARG | 383 | C2 | C | 101 | 3.5 |
| cIDR | NE | ARG | 383 | O2 | C | 101 | 2.8 |
| cIDR | NE | ARG | 383 | N3 | C | 101 | 3.6 |
| cIDR | CZ | ARG | 383 | C2 | C | 101 | 4.2 |
| cIDR | CZ | ARG | 383 | O2 | C | 101 | 3.3 |
| cIDR | CZ | ARG | 383 | N3 | C | 101 | 4.3 |
| cIDR | NH2 | ARG | 383 | C2 | C | 101 | 3.9 |
| cIDR | NH2 | ARG | 383 | O2 | C | 101 | 2.8 |

| IDR name | Protein atom name | Residue name | Residue number | RNA atom name | Nucleotide | Nucleotide number | Distance |
|---|---|---|---|---|---|---|---|
| cIDR | NH2 | ARG | 383 | N3 | C | 101 | 4.3 |
| cIDR | NH2 | ARG | 383 | N6 | A | 102 | 4.2 |
| cIDR | C | GLY | 384 | O2' | C | 99 | 4.3 |
| cIDR | C | GLY | 384 | O2 | C | 99 | 4.1 |
| cIDR | C | GLY | 384 | N2 | G | 109 | 3.9 |
| cIDR | O | GLY | 384 | O2' | C | 99 | 3.8 |
| cIDR | O | GLY | 384 | C1' | C | 99 | 4.4 |
| cIDR | O | GLY | 384 | C2 | C | 99 | 4.3 |
| cIDR | O | GLY | 384 | O2 | C | 99 | 3.2 |
| cIDR | O | GLY | 384 | O2' | C | 99 | 4.3 |
| cIDR | O | GLY | 384 | C2 | G | 109 | 4.4 |
| cIDR | O | GLY | 384 | N2 | G | 109 | 3.1 |
| cIDR | N | GLY | 385 | O2' | C | 99 | 4.4 |
| cIDR | N | GLY | 385 | O2 | C | 99 | 4.2 |
| cIDR | CA | GLY | 385 | O4' | C | 99 | 4.3 |
| cIDR | CA | GLY | 385 | O2' | C | 99 | 4 |
| cIDR | CA | GLY | 385 | C1' | C | 99 | 3.8 |
| cIDR | CA | GLY | 385 | C2 | C | 99 | 4.4 |
| cIDR | CA | GLY | 385 | O2 | C | 99 | 3.4 |
| cIDR | CA | GLY | 385 | N2 | G | 110 | 4.2 |
| cIDR | C | GLY | 385 | N2 | G | 110 | 4.2 |
| cIDR | N | ARG | 386 | C2 | G | 110 | 4.4 |
| cIDR | N | ARG | 386 | N2 | G | 110 | 3.3 |
| cIDR | CA | ARG | 386 | N2 | G | 110 | 4 |
| cIDR | C | ARG | 386 | O2' | G | 110 | 4.3 |
| cIDR | C | ARG | 386 | C4' | A | 111 | 4.4 |

| IDR name | Protein atom name | Residue name | Residue number | RNA atom name | Nucleotide | Nucleotide number | Distance |
|---|---|---|---|---|---|---|---|
| cIDR | C | ARG | 386 | O4' | A | 111 | 3.8 |
| cIDR | C | ARG | 386 | O2' | A | 111 | 4.2 |
| cIDR | C | ARG | 386 | C1' | A | 111 | 4.1 |
| cIDR | O | ARG | 386 | O2' | G | 110 | 4.1 |
| cIDR | O | ARG | 386 | N2 | G | 110 | 4.2 |
| cIDR | O | ARG | 386 | N3 | G | 110 | 4.3 |
| cIDR | O | ARG | 386 | O2' | G | 110 | 3.1 |
| cIDR | O | ARG | 386 | C4' | A | 111 | 4 |
| cIDR | O | ARG | 386 | O4' | A | 111 | 3.3 |
| cIDR | O | ARG | 386 | C1' | A | 111 | 3.9 |
| cIDR | CB | ARG | 386 | O2 | C | 98 | 4.2 |
| cIDR | CB | ARG | 386 | N2 | G | 110 | 3.6 |
| cIDR | CB | ARG | 386 | O2' | A | 111 | 4.1 |
| cIDR | CB | ARG | 386 | C1' | A | 111 | 4.1 |
| cIDR | CB | ARG | 386 | N3 | A | 111 | 4.4 |
| cIDR | CG | ARG | 386 | O2' | C | 98 | 4 |
| cIDR | CG | ARG | 386 | O2 | C | 98 | 3.7 |
| cIDR | CG | ARG | 386 | N2 | G | 110 | 4 |
| cIDR | CD | ARG | 386 | O2' | C | 98 | 3.9 |
| cIDR | CD | ARG | 386 | C1' | C | 98 | 3.8 |
| cIDR | CD | ARG | 386 | C2 | C | 98 | 4.4 |
| cIDR | CD | ARG | 386 | O2 | C | 98 | 3.5 |
| cIDR | CD | ARG | 386 | N3 | A | 111 | 4.4 |
| cIDR | CZ | ARG | 386 | O2' | A | 111 | 3.7 |
| cIDR | CZ | ARG | 386 | N3 | A | 111 | 4.2 |
| cIDR | CZ | ARG | 386 | O2' | A | 111 | 4 |

| IDR name | Protein atom name | Residue name | Residue number | RNA atom name | Nucleotide | Nucleotide number | Distance |
|---|---|---|---|---|---|---|---|
| cIDR | CZ | ARG | 386 | C4' | A | 112 | 4.4 |
| cIDR | CZ | ARG | 386 | O4' | A | 112 | 3.3 |
| cIDR | CZ | ARG | 386 | C1' | A | 112 | 3.9 |
| cIDR | NH1 | ARG | 386 | C2' | A | 111 | 3.6 |
| cIDR | NH1 | ARG | 386 | O2' | A | 111 | 3.1 |
| cIDR | NH1 | ARG | 386 | C1' | A | 111 | 3.8 |
| cIDR | NH1 | ARG | 386 | N9 | A | 111 | 4.4 |
| cIDR | NH1 | ARG | 386 | C2 | A | 111 | 3.7 |
| cIDR | NH1 | ARG | 386 | N3 | A | 111 | 3 |
| cIDR | NH1 | ARG | 386 | C4 | A | 111 | 4 |
| cIDR | NH1 | ARG | 386 | O2' | A | 111 | 3.5 |
| cIDR | NH1 | ARG | 386 | C4' | A | 112 | 4.2 |
| cIDR | NH1 | ARG | 386 | O4' | A | 112 | 2.9 |
| cIDR | NH1 | ARG | 386 | C1' | A | 112 | 3.4 |
| cIDR | NH1 | ARG | 386 | N9 | A | 112 | 3.8 |
| cIDR | NH1 | ARG | 386 | C8 | A | 112 | 4.3 |
| cIDR | NH1 | ARG | 386 | C4 | A | 112 | 4.4 |
| cIDR | NH2 | ARG | 386 | O2' | A | 111 | 3.9 |
| cIDR | NH2 | ARG | 386 | O2' | A | 111 | 3.8 |
| cIDR | NH2 | ARG | 386 | C5' | A | 112 | 4.4 |
| cIDR | NH2 | ARG | 386 | C4' | A | 112 | 3.6 |
| cIDR | NH2 | ARG | 386 | O4' | A | 112 | 2.9 |
| cIDR | NH2 | ARG | 386 | O2' | A | 112 | 4.2 |
| cIDR | NH2 | ARG | 386 | C1' | A | 112 | 3.6 |
| cIDR | N | GLY | 387 | C4' | A | 111 | 4.3 |
| cIDR | N | GLY | 387 | O4' | A | 111 | 4.2 |

| IDR name | Protein atom name | Residue name | Residue number | RNA atom name | Nucleotide | Nucleotide number | Distance |
|---|---|---|---|---|---|---|---|
| cIDR | N | GLY | 387 | O2' | A | 111 | 3.9 |
| cIDR | N | GLY | 387 | O2' | A | 111 | 4.4 |
| cIDR | CA | GLY | 387 | C4' | A | 111 | 3.8 |
| cIDR | CA | GLY | 387 | O4' | A | 111 | 4.1 |
| cIDR | CA | GLY | 387 | O2' | A | 111 | 4 |
| cIDR | CA | GLY | 387 | O2' | A | 111 | 4.4 |
| cIDR | C | GLY | 387 | C4' | A | 111 | 3.7 |
| cIDR | C | GLY | 387 | O4' | A | 111 | 4.4 |
| cIDR | C | GLY | 387 | C3' | A | 111 | 4.3 |
| cIDR | C | GLY | 387 | O3' | A | 111 | 4 |
| cIDR | C | GLY | 387 | C2' | A | 111 | 4.4 |
| cIDR | C | GLY | 387 | O2' | A | 111 | 3.4 |
| cIDR | C | GLY | 387 | O2' | A | 111 | 3.5 |
| cIDR | O | GLY | 387 | C4' | A | 111 | 4.3 |
| cIDR | O | GLY | 387 | O3' | A | 111 | 4.1 |
| cIDR | O | GLY | 387 | C2' | A | 111 | 4.2 |
| cIDR | O | GLY | 387 | O2' | A | 111 | 3 |
| cIDR | O | GLY | 387 | O2' | A | 111 | 2.9 |
| cIDR | O | GLY | 387 | C5' | A | 112 | 4.4 |
| cIDR | N | ARG | 388 | C5' | A | 111 | 4.3 |
| cIDR | N | ARG | 388 | C4' | A | 111 | 3.8 |
| cIDR | N | ARG | 388 | C3' | A | 111 | 4.3 |
| cIDR | N | ARG | 388 | O3' | A | 111 | 3.7 |
| cIDR | N | ARG | 388 | O2' | A | 111 | 4.1 |
| cIDR | N | ARG | 388 | O2' | A | 111 | 3.9 |
| cIDR | CA | ARG | 388 | C4' | A | 111 | 4.4 |

| IDR name | Protein atom name | Residue name | Residue number | RNA atom name | Nucleotide | Nucleotide number | Distance |
|---|---|---|---|---|---|---|---|
| cIDR | CA | ARG | 388 | O3' | A | 111 | 3.4 |
| cIDR | CA | ARG | 388 | O2' | A | 111 | 4.4 |
| cIDR | CA | ARG | 388 | O2' | A | 111 | 3.9 |
| cIDR | CA | ARG | 388 | P | A | 112 | 4.2 |
| cIDR | CA | ARG | 388 | OP1 | A | 112 | 3.7 |
| cIDR | CB | ARG | 388 | O3' | A | 111 | 3.9 |
| cIDR | CB | ARG | 388 | OP1 | A | 112 | 3.8 |
| cIDR | CG | ARG | 388 | C5' | A | 111 | 3.8 |
| cIDR | CG | ARG | 388 | C4' | A | 111 | 4.1 |
| cIDR | CG | ARG | 388 | C3' | A | 111 | 4.3 |
| cIDR | CG | ARG | 388 | O3' | A | 111 | 3.3 |
| cIDR | CG | ARG | 388 | P | A | 112 | 4.1 |
| cIDR | CG | ARG | 388 | OP1 | A | 112 | 3.6 |
| cIDR | NH1 | ARG | 388 | OP1 | A | 111 | 3.3 |
| cIDR | NH1 | ARG | 388 | C5' | A | 111 | 4.2 |
| cIDR | N | GLY | 389 | OP1 | A | 112 | 4.2 |
| cIDR | N | GLY | 389 | C5' | A | 112 | 4.3 |
| cIDR | O | GLY | 390 | C4' | A | 112 | 4.3 |

Supplementary Table S2: Intermolecular H-bonds formed in the two replicates of FUS RRM bound to U1 snRNA.

| Residues | RNA nucleotides | H-bond occupancy (%) | |
|---|---|---|---|
| | | Traj1 | Traj2 |
| PHE265 | G91 | 14.44 | |
| ARG269 | A111 | | 10.38 |
| ASP270 | C118 | 19.84 | |
| ASP283 | C117 | 10.03 | |
| ASN285 | U107 | | 22.79 |
| LYS312 | U95 | 33.47 | |
| | A94 | 14.01 | |
| | G93 | | 10.12 |
| LYS315 | U97 | 10.62 | |
| | U96 | 18.94 | |
| | A94 | | 10.45 |
| LYS316 | G108 | 11.60 | |
| TYR325 | U107 | 26.56 | |
| THR326 | U105 | 30.14 | 46.67 |
| ARG328 | U105 | | 15.14 |
| | A104 | 12.59 | 11.82 |
| LYS334 | U105 | 11.55 | |
| PHE368 | G106 | 27.84 | |
| THR370 | G106 | | 21.26 |
| ARG371 | U107 | 35.08 | 23.95 |
| | G109 | 27.47 | 14.01 |
| ARG372 | G106 | | 40.06 |
| | U105 | | 19.93 |
| ALA373 | G106 | | 39.42 |

| | | | |
|---|---|---|---|
| ASN376 | U105 | | 30.33 |
| ARG377 | U105 | 40.50 | |
| GLY380 | G110 | 11.45 | |
| ASN381 | C100 | 13.52 | |
| | G108 | | 13.13, 22.10, 45.79 |
| | C101 | | 18.48 |
| ARG383 | C101 | 19.46 | |
| | G109 | 37.46 | |
| | A103 | | 29.79 |
| GLY384 | G109 | | 19.24 |
| GLY385 | G110 | 10.48 | |
| ARG386 | C99 | | 11.32 |
